# Human milk components interact with infant genomics to modulate gut microbiota, childhood asthma and atopy

**DOI:** 10.1101/2025.09.19.676891

**Authors:** Sara A. Stickley, Zhi Yi Fang, Amirthagowri Ambalavanan, Elizabeth George, Charisse Petersen, Darlene L.Y. Dai, Shirin Moossavi, Kozeta Miliku, Lars Bode, Catherine J. Field, Piushkumar J. Mandhane, Elinor Simons, Theo J. Moraes, Michael G. Surette, Meghan B. Azad, Stuart E. Turvey, Padmaja Subbarao, Qingling Duan

## Abstract

The benefits of breastfeeding are well established; however, the mechanisms by which human milk components (HMCs) impact children’s long-term health remain poorly understood. We leveraged datasets from the CHILD Cohort Study to explore how exposure to variable HMCs— including oligosaccharides (HMOs), fatty acids (HMFAs), and microbiota (HMM)—may influence infants’ gut microbiota and risk of childhood asthma and atopy. We identified HMCs (e.g., HMO lacto-N-fucopentaose III and HMFA linoleic acid) associated with gut microbes and microbial networks implicated in atopic diseases. Additionally, we determined that HMCs (e.g., HMM *Pseudomonas oryzihabitans)* interact with infants’ polygenic risk scores (PRSs) to influence these gut microbial features. Integration of HMCs, gut microbiota, and disease-associated PRSs into an unsupervised machine-learning model that clustered two groups of infants with differing disease prevalence. Our findings suggest that HMCs influence childhood asthma and atopy through modifications to the gut microbiota and modulated by interactions with infant genomics.

## Introduction

Breastfeeding or human milk is known to have long-term benefits for infants, including supporting immune system functioning, neurodevelopment, and protection from childhood obesity, respiratory and other infections^1–6^. However, past investigations reported inconsistent associations between breastfeeding and childhood allergic diseases^7–13^, which may be due in part to the consideration of breastfeeding as a single homogeneous exposure. Instead, human milk is a complex and heterogeneous exposure as it is composed of thousands of bioactive components, such as human milk oligosaccharides (HMOs), fatty acids (HMFAs), microbes (HMM), proteins, and hormones^14–18^. Additionally, human milk components (HMCs) have been shown to vary among individuals due to differences in genomics or environmental exposures^19–29^.

In recent years, research has shifted to focus on how variable exposure to specific HMCs may impact the health of human milk-fed infants^19,20,29–32^. For example, previous work from our team reported that HMOs and HMM modulate respiratory and atopic health of milk-fed infants^19,20^. However, further research is needed to explore the underlying mechanisms through which these and other HMCs may influence childhood health and disease. One potential mechanism through which human milk exposure may impact childhood health is via the early-life gut microbiota, as it has been proposed that HMCs may act as prebiotics or probiotics, thereby shaping infant gut microbial colonization^33–35^.

Previous studies have reported that both breastfeeding practices^36–38^ and variations in HMCs are associated with infant gut microbial composition^24,38–44^. A recent investigation from the CHILD Cohort Study determined that breastfeeding duration may impact the colonization of both the infant nasal and gut microbiota by age 1 year, and in turn, impact childhood asthma susceptibility^45^. Additional studies are necessary to further elucidate how specific HMCs may impact the infant gut microbiota and influence childhood health. Moreover, the role of infant genomics in modulating the impact of HMCs on the early-life gut microbiota remains poorly understood.

Our study provides novel insights into how early-life exposure to variable HMOs, HMFAs, and HMM may shape the infant gut microbiota in the first year of life, influencing the long-term risk of childhood asthma and atopy. We also examine how exposure to HMCs may shape the gut microbiota differently depending on the children’s genetic susceptibility to these diseases. Furthermore, we utilize an unsupervised multi-omics subject network approach, known as similarity network fusion^46^, to integrate HMCs, gut microbiota and infant genomics to identify infants with increased risk of childhood asthma and atopy. Overall, our findings indicate that exposure to variable HMCs may influence susceptibility to childhood asthma and atopy, possibly through changes to the infant gut microbiota and depending on interactions with infant genomics. A better understanding of the mechanisms by which breastfeeding or human milk impact childhood health may facilitate early-life intervention strategies, such as dietary supplementation or substitution, particularly for children with high genetic risk of chronic diseases or those not receiving human milk.

## Results

### Study overview and participants

Our study leveraged datasets from the Canadian CHILD Cohort Study, which follows 3,542 infants born from singleton pregnancies between 2008–2012 and their families (**Figure 1**). This investigation focused on subsets of mother-infant dyads with maternal HMOs, HMFAs, and HMM quantified from human milk samples collected at 3-4 months postpartum and infant gut microbiota profiles determined from stool samples collected at 3 months or 1 year of age (**Figure 1, Table S1, Table S2**). To explore the potential impact of HMCs on the infant gut microbiota, we focused on gut microbial amplicon sequence variants (ASVs) and network clusters of co-occurring microbes that we previously associated with asthma and atopy prevalence^47^. These gut microbial features included: 2 ASVs and 4 network clusters of co-occurring microbes in the infant gut at age 3 months; and 3 ASVs and 7 microbial clusters in the infant gut at age 1 year^47^ (**Table S3, Table S4, Table S5**). We referred to the microbial features previously correlated with increased asthma/atopy risk (*Clostridium sensu stricto sp. 4, Enterobacter cloacae,* and *Megasphaera micronuciformis,* and *black, brown, green, red,* and *yellow* clusters) as disease-elevated, and microbial features previously correlated with decreased asthma/atopy risk (*Fusicatenibacter sp. 3* and *Agathobacter sp. 6,* and *blue, turquoise, orange, magenta, salmon,* and *purple* clusters) as disease-reduced (**Table S3**). To study how host genomic predisposition may modulate the effect of HMCs on the infant gut microbiota, we focused on a further subset of infants with genomic profiles (**Figure 1**). Lastly, we integrated all HMCs, gut microbiota, and genomic profiles in a similarity network fusion model, which is an unsupervised machine-learning approach, to group infants with different prevalence of childhood asthma and atopy. We explored asthma and atopy outcomes including: asthma diagnosis at age 5 years, recurrent wheeze between ages 2 to 5 years, and atopic dermatitis, food sensitization, and inhalant sensitization between ages 3 to 5 years (**Table S6**). A graphical overview of our study design is shown in **Figure S1**.

**Figure 1.**
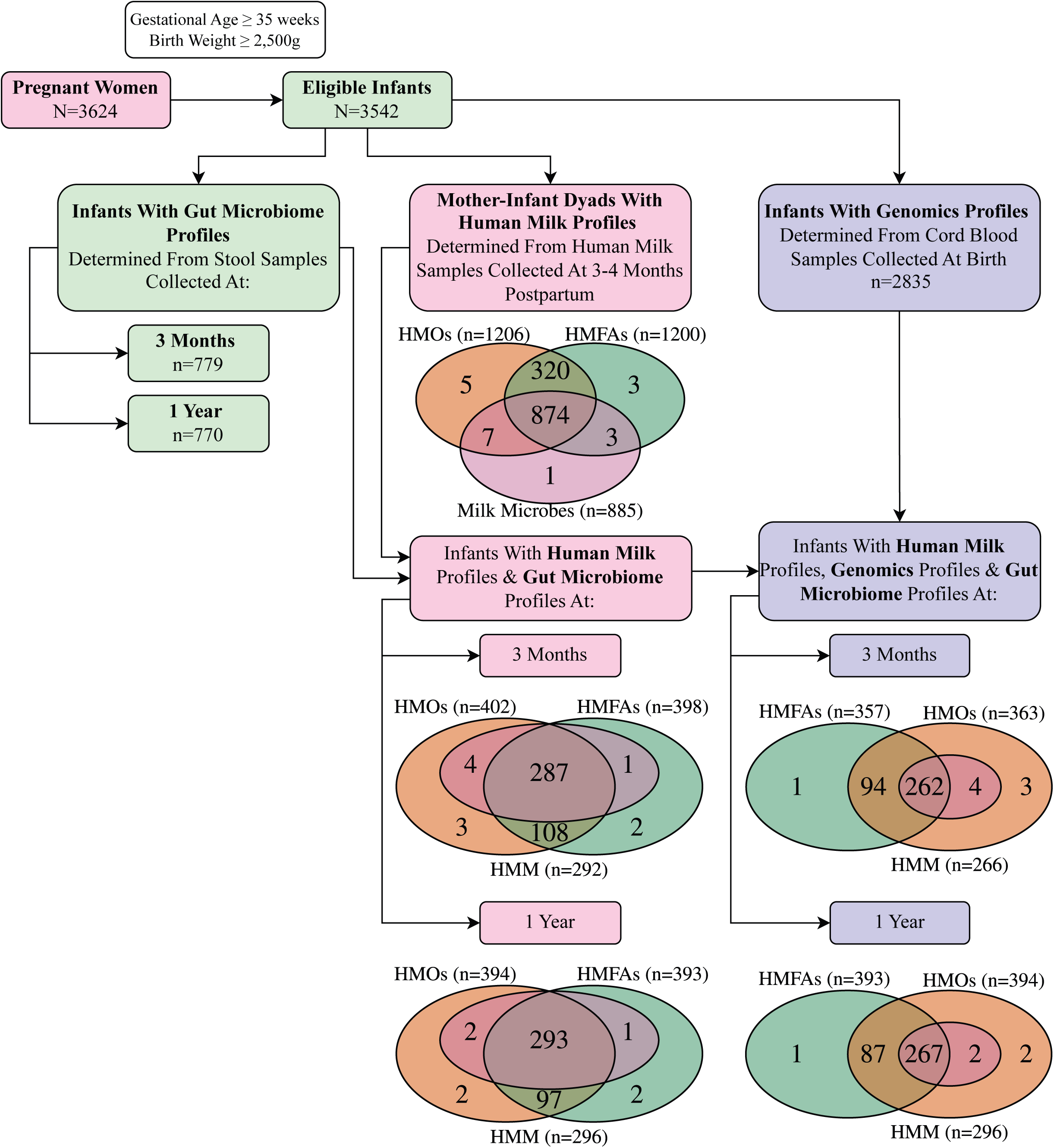
Study overview. **(A)** Timeline of biological data and demographic information collected in the CHILD Cohort Study from gestation to age 5 years. **(B)** A subset of infants had both HMC and gut microbiome data at 3 months and 1 year of age. Smaller subsets also had genomics data.

### Exposure to HMCs was associated with the infant gut microbiota

We identified that early-life exposures to HMOs, HMFAs, or HMM were associated with gut ASVs and network clusters of correlated microbiota at 3 months or 1 year of age, which were previously correlated with childhood asthma and atopy (**Figure 2, Table S7**). Our analysis identified 3 gut ASVs correlated with HMFA exposure (**Figure 2A**). For example, PUFA measures (e.g., linoleic acid (LA, 18:2n6), P_Bonferroni_=0.02, β=0.4) were associated with disease-elevated *Clostridium sensu stricto sp. 4* abundance at 3 months, suggesting harmful effects. Additionally, MUFAs oleic acid (18:1n9) and nervonic acid (24:1n9) were associated with disease-reduced *Agathobacter sp. 6* (P_Bonferroni_=0.02, β=0.5) and *Fusicatenibacter sp. 3* (P_Bonferroni_=0.04, β=0.3) abundances at 1 year, respectively, suggesting protective effects.

**Figure 2.**
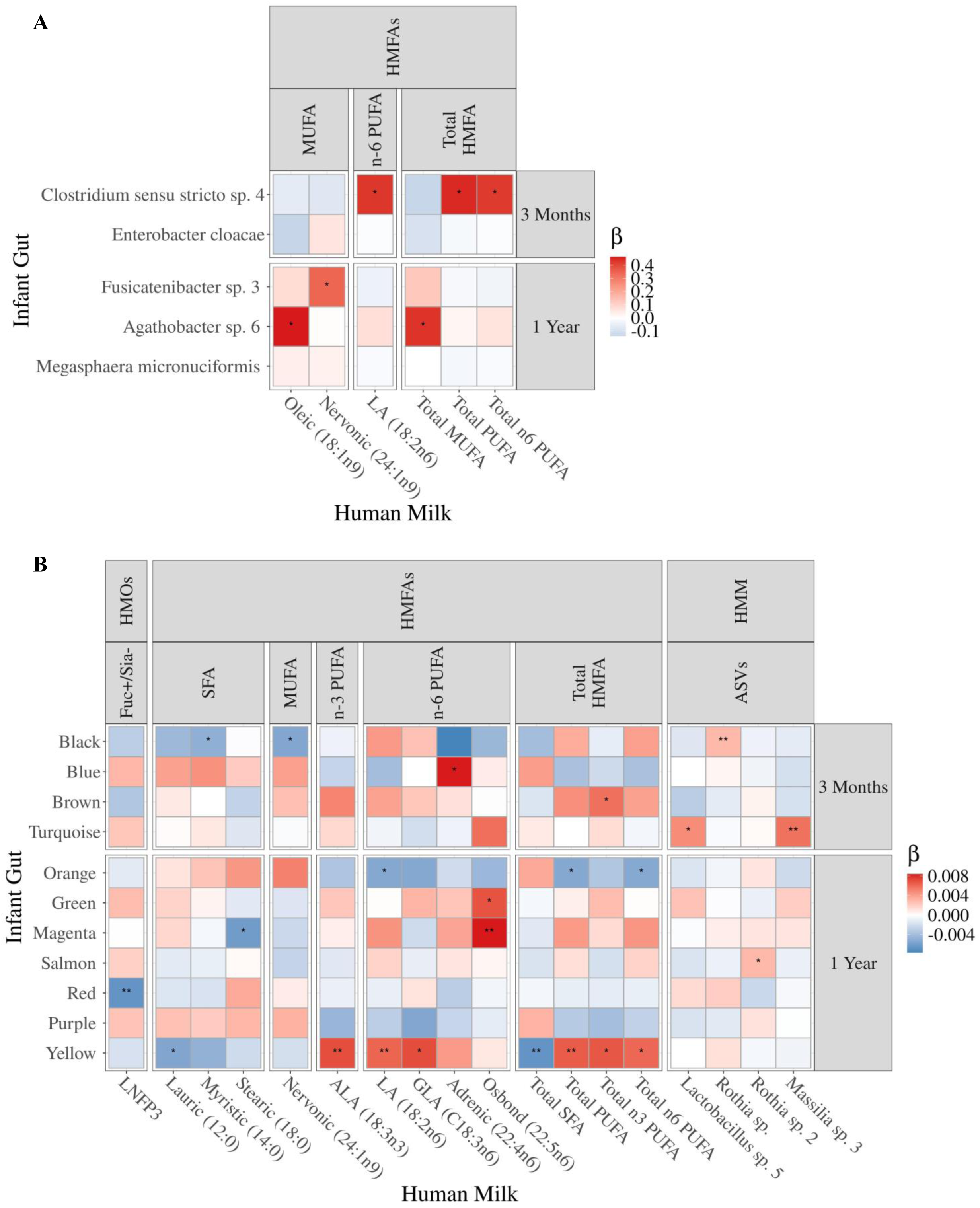
Early life gut microbiota were associated with exposure to variable HMCs. The heatmaps show significant associations between HMOs, HMFAs, and HMM and the asthma and atopy-implicated^47^ **(A)** gut amplicon sequence variants (ASV) abundances and **(B)** gut microbial network clusters at 3 months or 1 year of age. Stars correspond to Bonferroni-adjusted P values as follows: * P < 0.05, ** P < 0.01, and *** P < 0.001. Using the Bonferroni correction, P values for gut ASV abundances were adjusted for the number of independent HMCs multiplied by the number of gut ASVs, while P values for gut microbial network clusters were adjusted for the number of independent HMCs. Beta estimates (β) indicate directionality of associations, where a positive β (red) indicates that the HMC was associated with an increased microbial feature, while negative β (blue) indicates that the HMC was associated with a decreased microbial feature.

We also determined that HMCs were associated with gut microbial network clusters (**Figure 2B**). For example, exposure to the HMO lacto-*N*-fucopentaose 3 (LNFP3) was correlated with lower abundance of the disease-elevated *red* gut microbial cluster at 1 year (P_Bonferroni_=8.6E-3, β=-6.7E-3), suggesting protective effects. Additionally, 14 HMFAs were associated with gut microbial clusters. For example, polyunsaturated fatty acid (PUFA) measures were associated with disease-elevated *brown* gut cluster at 3 months (e.g., total n-3 PUFAs, P_Bonferroni_=0.02, β=5.8E-3) and the disease-elevated *yellow* gut cluster at 1 year (e.g., α-linolenic acid (ALA) (18:3n3), P_Bonferroni_=5.7E-3, β=7.1E-3), suggesting harmful effects. In contrast, the PUFAs, osbond acid (22:5n6), was associated with the disease-reduced *magenta* gut cluster (P_Bonferroni_=7.9E-3, β=8.4E-3) at 1 year, suggesting protective effects. Lastly, 4 HMM ASVs were associated with gut clusters. HMM *Rothia sp.* was correlated with the disease-elevated *black* gut cluster at 3 months (P_Bonferroni_=1.7E-3, β=2.9E-3), suggesting harmful effects, while *Massilia sp. 3* (P_Bonferroni_=1.4E-3, β=5.8E-3) and *Lactobacillus sp. 5* (P_Bonferroni_=0.04, β=4.7E-3) were associated with the disease-reduced *turquoise* cluster, suggesting protective effects. In contrast, a different HMM *Rothia* species, *Rothia sp. 2,* was associated with the disease-reduced *salmon* gut cluster at 1 year (P_Bonferroni_=0.04, β=2.9E-3), suggesting protective effects.

### Asthma and atopy PRSs were associated with disease prevalence

In order to quantify infants’ polygenic risk of developing asthma or atopy during childhood, we computed two polygenic risk scores (PRSs)–one for asthma and one for atopy–based on 1,107,307 and 1,097,013 SNPs previously associated with asthma^48^ and atopy^49^, respectively. We correlated these PRSs with childhood asthma and atopic outcomes as shown in **Figure 3A** and **Figure 3B**. Increased asthma PRS (**Figure 3A)** was correlated with asthma diagnosed by age 5 years (P=2.5E-5, β=0.4) and recurrent wheeze episodes between ages 2–5 years (P=6.1E-9, β=0.4). Increased atopy PRS (**Figure 3B**) was correlated with prevalence of AD (P=6.1E-7, β=0.4), food sensitization (P=1.3E-5, β=0.5), and inhalant sensitization (P=3.0E-7, β=0.3). PRSs were z-score normalized and categorized as low (< −1), moderate (−1 to 1), and high (> 1) for visualization purposes.

**Figure 3.**
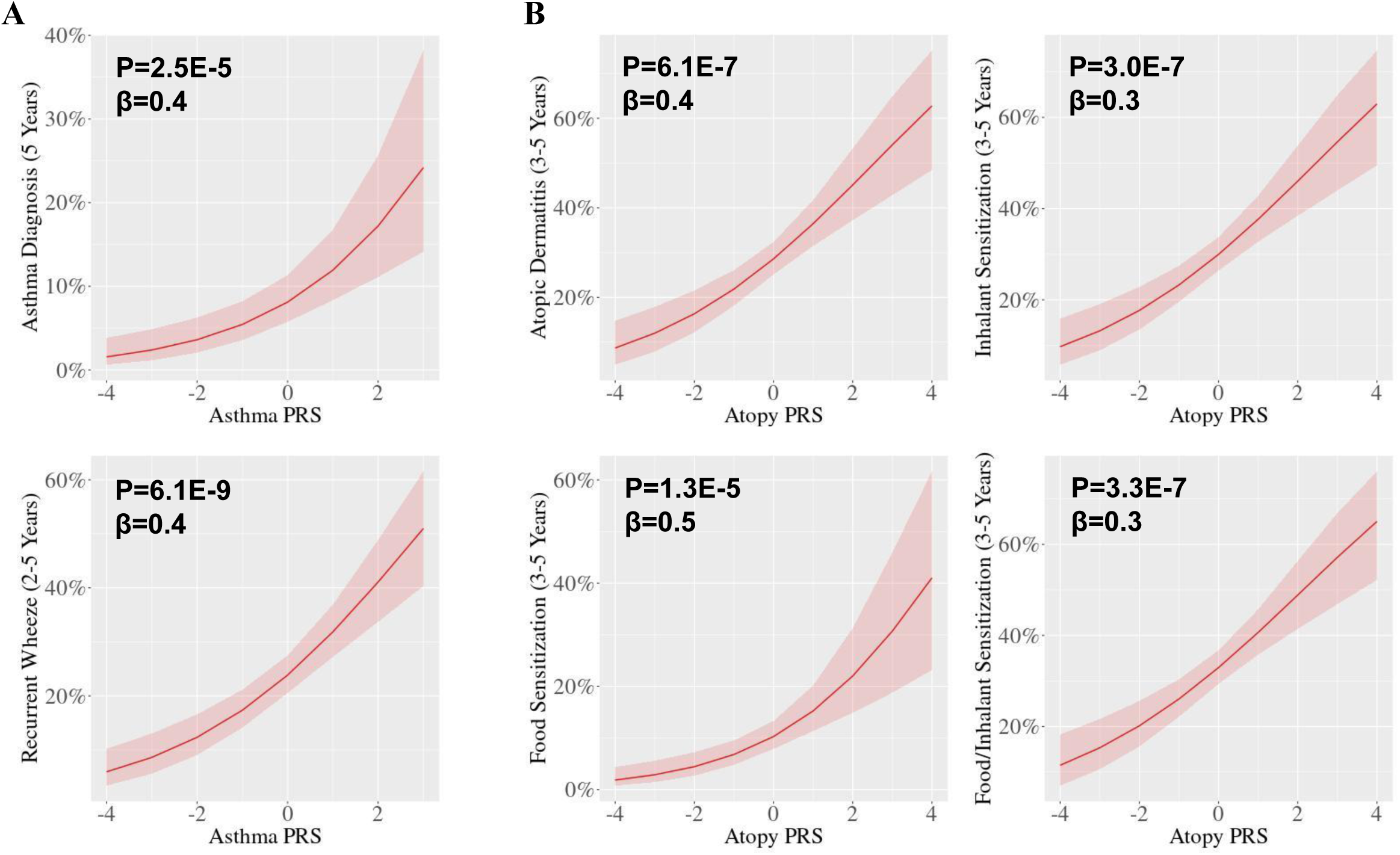
Infant polygenic risk scores (PRSs) were associated with asthma and atopy outcomes. **(A)** Increased asthma PRS was associated with increased prevalence of physician-diagnosed asthma at age 5 years and childhood recurrent wheeze between ages 2–5 years, as reported via questionnaires. **(B)** Increased atopy PRS was associated with increased prevalence of physician-diagnosed AD and sensitization to food and/or inhalant allergens as determined by skin-prick testing (SPT) between ages 3–5 years.

### Interactions between PRSs and HMCs were associated with composition of infant gut microbiota

We conducted gene-by-environment (G×E) interaction analysis to determine whether the associations between HMCs, such as HMOs, HMFAs and HMM (E), and the gut microbiota are dependent on the infants’ genomic predisposition (PRSs or G) for asthma and atopy (**Figure 4** and **Figure 5, Table S8** and **Table S9**).

**Figure 4.**
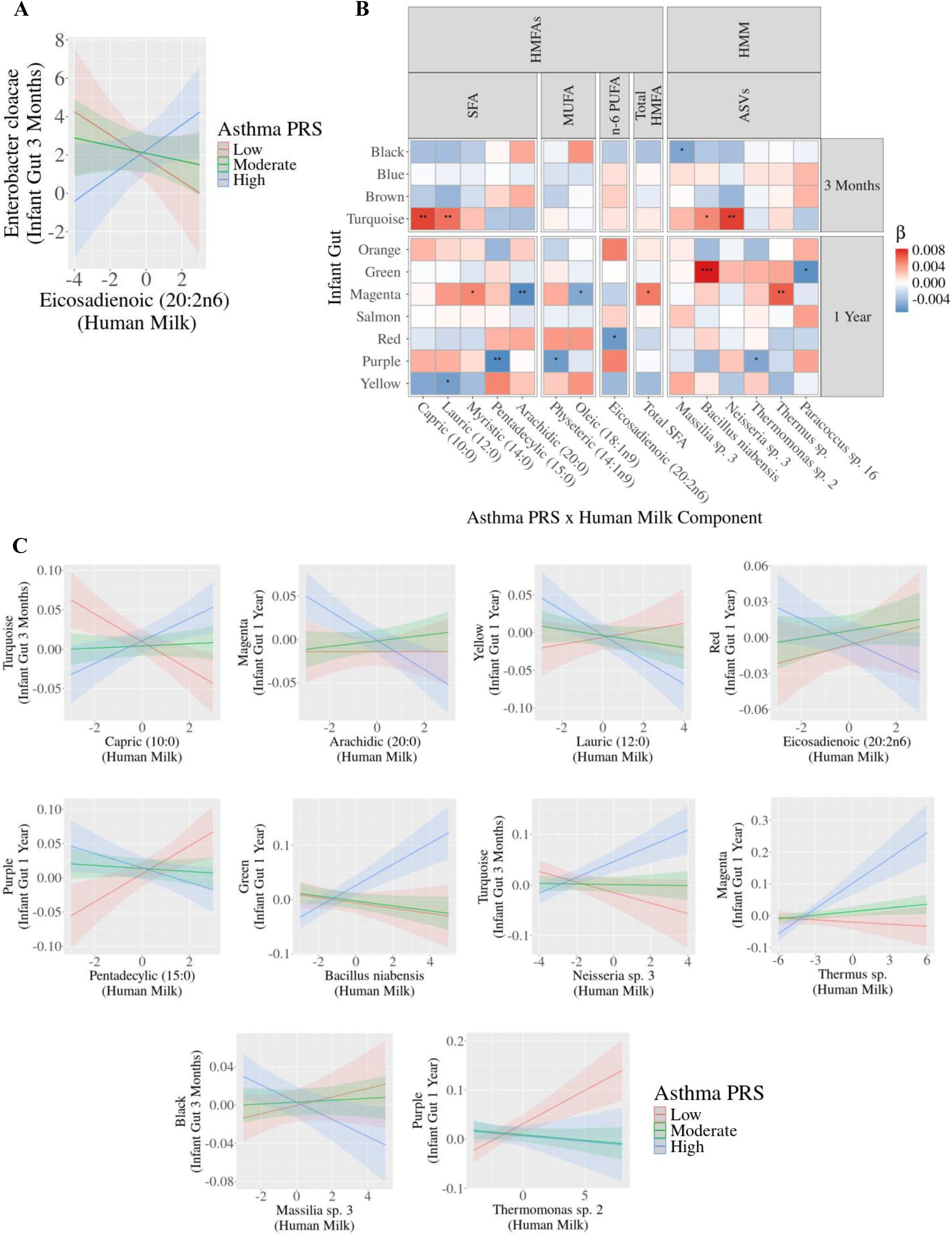
Interactions between asthma PRS and HMCs were associated with gut microbiota. **(A)** The association between *Enterobacter cloacae* in the gut at 3 months and the HMFA eicosadienoic acid (20:2n6) (P<0.05) was dependent on the infants’ asthma PRS. **(B)** The heatmap shows that gut microbial network clusters were significantly associated with interactions between asthma PRS and HMCs (HMOs, HMFAs, and HMM). Stars correspond to Bonferroni-adjusted P values: * P < 0.05, ** P < 0.01, and *** P < 0.001. For gut ASVs, we corrected for the number of independent HMCs multiplied by the number of gut ASVs tested. For gut microbial network clusters, we adjusted for the number of independent HMCs. Beta estimates (β) indicate directionality of associations among infants with higher asthma PRS, where a positive β (red) indicates that the HMC was associated with increased microbial feature, while negative β (blue) indicates that the HMC was associated with decreased microbial feature. **(C)** Additional interaction plots show examples of significant interactions between asthma PRS and HMCs associated with gut microbial network clusters. In **(A)** and **(C),** the x-axes indicate the normalized HMC, and the y-axes indicate gut microbial features–either normalized ASV abundances or microbial network cluster eigenvalues. The legend corresponds to the Z-score-normalized asthma PRSs which was categorized into low (PRS < −1), moderate (PRS of −1 to 1), and high (PRS > 1) for plotting.

**Figure 5.**
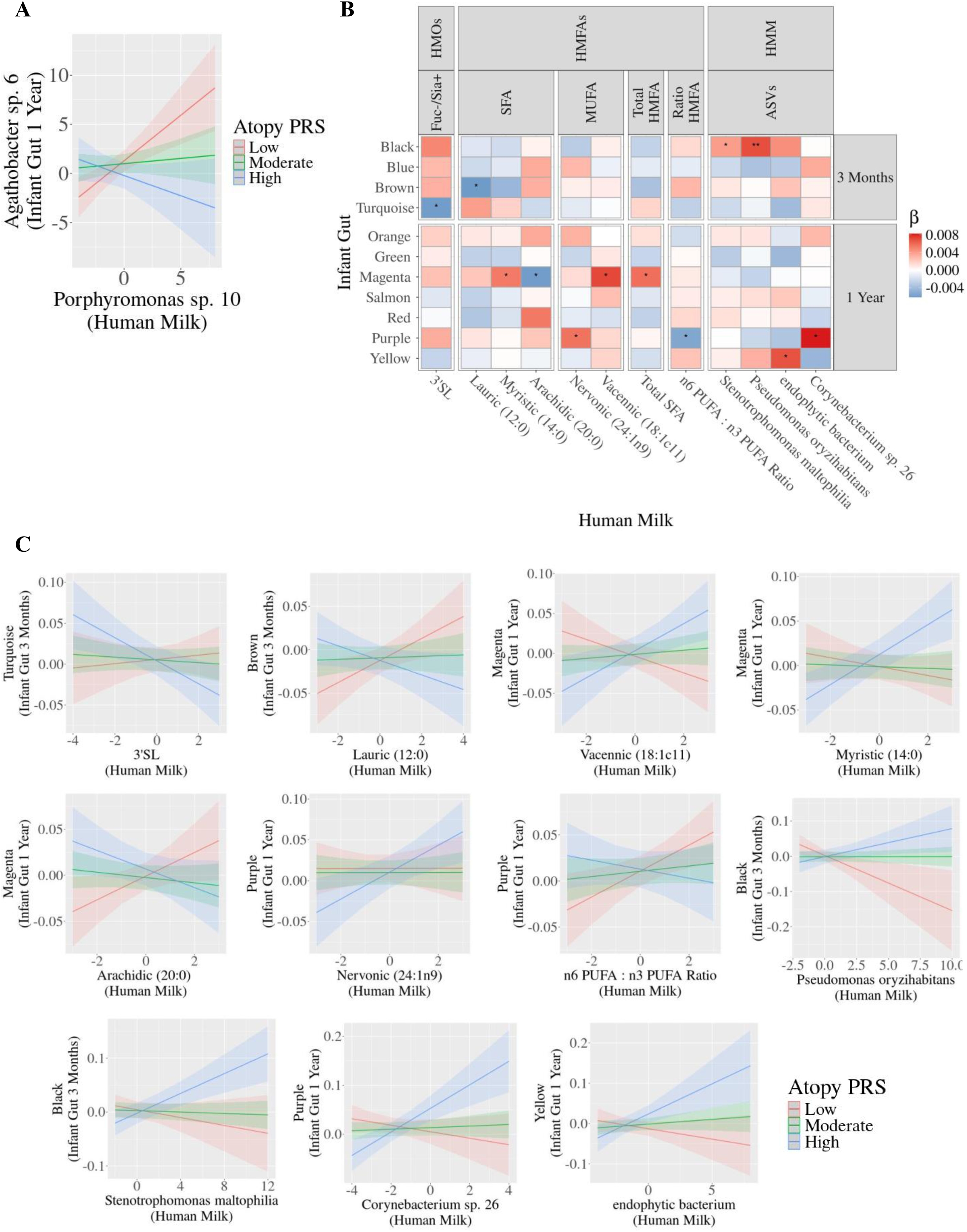
Interactions between atopy PRS and HMCs were associated with gut microbiota. **(A)** The association between *Agathobacter sp. 6* in the gut at 1 year associated with the HMM ASV *Porphyromonas sp. 10* (P<0.05) was dependent on infants’ atopy PRS. **(B)** The heatmap shows that gut microbial network clusters were significantly associated with interactions between atopy PRS and HMCs (HMOs, HMFAs, and HMM). Stars correspond to Bonferroni-adjusted P values: * P < 0.05, ** P < 0.01, and *** P < 0.001. For gut ASVs, we corrected for the number of independent HMCs multiplied by the number of gut ASVs. For gut microbial network clusters, we adjusted for the number of independent HMCs. Beta estimates (β) indicate directionality of associations among infants with higher atopy PRS, where a positive β(red) indicates that the HMC was associated with increased microbial feature, while negative β(blue) indicates that the HMC was associated with decreased microbial feature. **(C)** Additional interaction plots show examples of significant interactions between atopy PRS and HMCs associated with gut microbial network clusters. In **(A)** and **(C),** the x-axes indicate the normalized HMC, and the y-axes indicate gut microbial features–either normalized ASV abundances or microbial network cluster eigenvalues. The legend corresponds to the Z-score-normalized atopy PRS which was categorized into low (PRS < −1), moderate (PRS of −1 to 1), and high (PRS > 1) for plotting.

#### Asthma PRS×HMCs

Among infants with high asthma PRS, those exposed to more HMFA eicosadienoic acid (20:2n6) had increased disease-elevated *E. cloacae* in the gut at 3 months (P_Bonferroni_=0.05, β=0.4), whereas those with lower PRS and the same exposure had decreased abundance of this gut ASV (**Figure 4A**).

We also identified 17 significant interactions between PRS and HMCs associated with gut microbial network clusters (**Figure 4B**, examples highlighted in **Figure 4C**). Interactions between asthma PRS and 9 HMFA measures were associated with gut clusters across both timepoints. For example, infants with high asthma PRS who were exposed to more SFAs (e.g., capric acid (10:0), P_Bonferroni_=8.4E-3, β=7.3E-3) had increased disease-reduced *turquoise* gut cluster at 3 months, whereas those with lower PRS and the same exposure had decreased *turquoise* gut cluster. Furthermore, infants with high asthma PRS and exposed to high arachidic acid (20:0) had decreased disease-reduced *magenta* gut cluster (P_Bonferroni_=3.2E-3, β=-7.0E-3). Similarly, infants with high asthma PRS exposed to more lauric acid (12:0) and eicosadienoic acid (20:2n6) had decreased disease-elevated *yellow* cluster (P_Bonferroni_=0.01, β=-5.9E-3) and *red* (P_Bonferroni_=0.01, β=-6.2E-3) clusters, respectively. In contrast to the infants with high asthma PRS, those with low asthma PRS and exposed to more pentadecylic acid (15:0) had increased disease-reduced *purple* gut cluster at 1 year (P_Bonferroni_=6.1E-3, β=-7.2E-3).

Additionally, our *<u>PRS×HMCs</u>* analyses identified interactions between asthma PRS and 6 HMM ASVs. Most significantly, infants with high PRS exposed to more HMM *Bacillus niabensis* had increased disease-elevated *green* gut cluster at 1 year (P_Bonferroni_=9.3E-4, β=8.2E-3). Also, infants with high PRS exposed to more HMM *Neisseria sp. 3* and *Thermus sp.* had increased disease-reduced *turquoise* gut cluster (P_Bonferroni_=4.2E-3, β=7.4E-3) at 3 months and *magenta* gut cluster (P_Bonferroni_=3.9E-3, β=6.4E-3) at 1 year, respectively. Moreover, infants with low PRS exposed to more HMM *Thermomonas sp. 2* had increased disease-reduced *purple* gut cluster at 1 year (P_Bonferroni_=0.03, β=-5.4E-3).

#### Atopy PRS×HMCs

Among infants with low atopy PRS, those exposed to the HMM *Porphyromonas sp. 10* had increased abundance of the disease-reduced *Agathobacter sp. 6* in the gut at 1 year (P_Bonferroni_=0.03, β=-0.5), whereas those with higher PRS and the same HMM exposure had a decreased abundance of this gut ASV (**Figure 5A**). Moreover, we identified 12 significant interactions between PRS and HMCs associated with gut microbial network clusters (**Figure 5B**, examples highlighted in **Figure 5C**). For example, infants with high atopy PRS exposed to higher concentrations of the HMO 3′-sialyllactose (3’SL) had a decreased abundance of the disease-reduced *turquoise* gut cluster at 3 months (P_Bonferroni_=0.03, β=-6.0E-3).

Additionally, we identified 7 *<u>PRS×HMFA</u>* interactions associated with gut clusters across both timepoints. Among infants with high atopy PRS, those exposed to more lauric acid (12:0) had decreased disease-elevated *brown* cluster (P_Bonferroni_=0.01, β=-6.2E-3), while those with lower PRS and the same HMFA exposure had increased abundance of this gut microbial cluster. Infants with high atopy PRS exposed to more SFAs or MUFAs (e.g., vaccenic acid (18:1c11), P_Bonferroni_=0.01, β=7.2E-3) also had increased disease-reduced *magenta* gut cluster at 1 year, while those with lower PRS exposed and the same HMFA exposure had decreased abundance of this gut cluster. In contrast, infants with lower PRS exposed to a higher n6 PUFA:n3 PUFA ratio had increased disease-reduced *purple* gut cluster at 1 year (P_Bonferroni_=0.04, β=-5.4E-3).

We identified 4 significant *<u>PRS×HMM</u>* ASV interactions. Among infants with high atopy PRS, those exposed to more HMM *Pseudomonas oryzihabitans* (P_Bonferroni_=7.8E-3, β=6.9E-3) and *Stenotrophomonas maltophilia* (P_Bonferroni_=0.02, β=4.0E-3) had increased disease-elevated *black* gut cluster at 3 months. In addition, infants with high atopy PRS and exposed to more HMM *Corynebacterium sp. 26* had increased disease-reduced *purple* gut cluster at 1 year (P_Bonferroni_=0.01, β=8.3E-3). In contrast, infants with lower atopy PRS exposed to any of these HMM ASVs had decreased gut microbial clusters.

### Multi-omics integration of HMCs, gut microbiota, and host PRSs clustered infants into groups associated with childhood asthma or atopy

The similarity network fusion (SNF) tool, a multi-omics subject network analysis approach, was used to identify groups of infants with similarities in human milk exposure, gut microbial composition, and host PRSs. Two multi-omics subject networks were constructed, which were referred to as: (1) the Human Milk-Gut Microbiota Network (**Figure S2**), integrating HMCs (19 HMOs, 28 HMFAs, and 185 HMM ASVs) and gut microbiota (125 gut ASVs at 3 months, 182 gut ASVs at 1 year) without host PRSs; and (2) the Human Milk-Gut Microbiota-PRS Network (**Figure 6**), integrating HMCs, gut microbiota, and host PRSs (associated with asthma and atopy). Subject similarity matrices were first constructed for each individual dataset, followed by integration (or fusion) into a multi-omics subject network. After network fusion, the eigen-gap and rotation cost algorithms were used to identify the ideal number of clusters for each constructed network. The optimal number of clusters was determined to be 2 for both networks. When infants were grouped using HMCs and gut microbiota without host PRSs, groups 1 and 2 had similar prevalence of asthma/atopy (**Figure S2B**). In contrast, when infants’ PRSs were incorporated into the model, group 1 had higher prevalence of asthma or atopy compared to group 2 (**Figure 6B)**.

**Figure 6.**
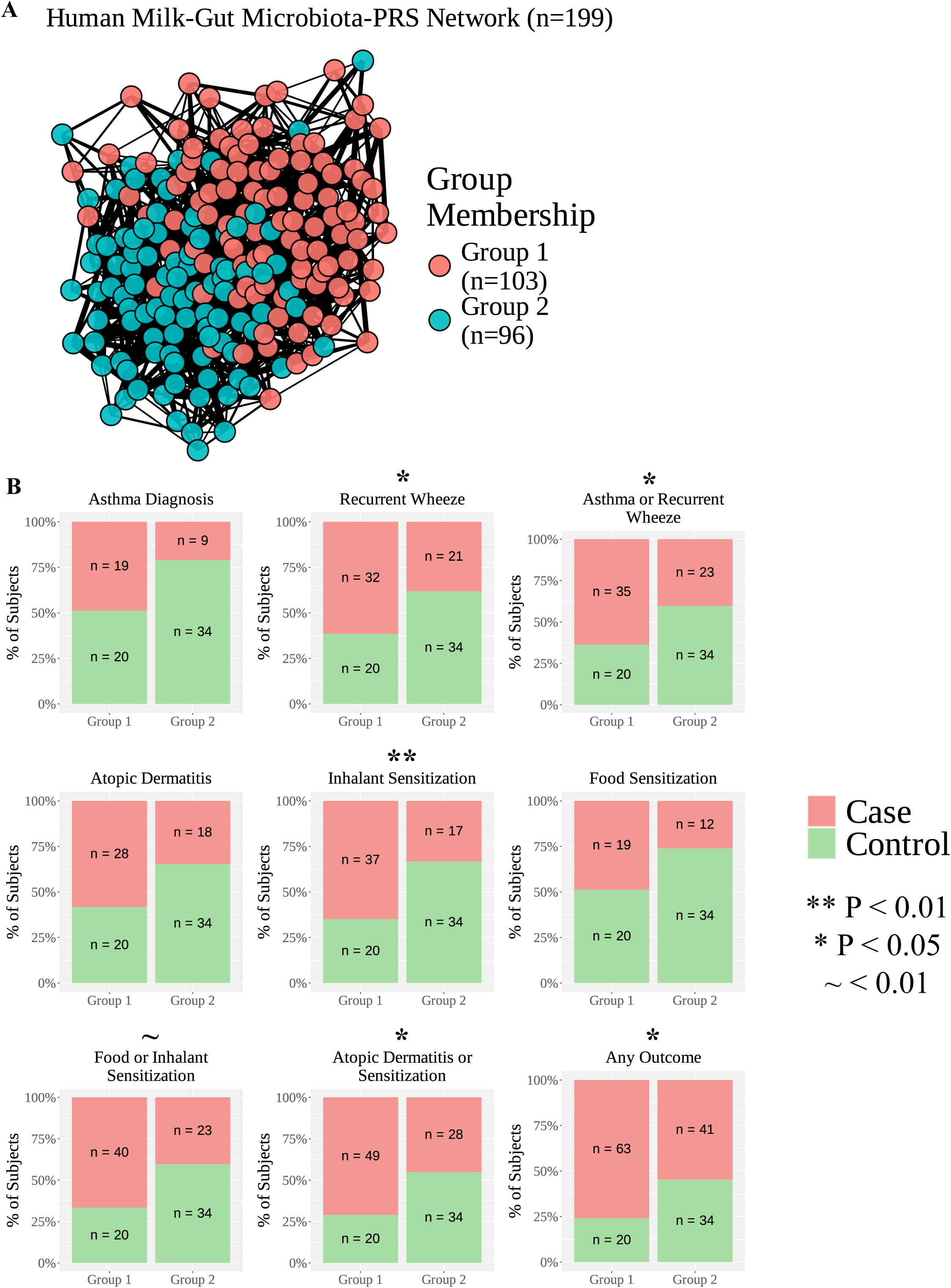
Multi-omics integration of HMCs, infant gut microbiota, and infant genomics identified groups of subjects associated with asthma and atopy risk. **(A)** Network plots show two infant groups identified from multi-omics subject network construction and clustering of 19 HMOs, 28 HMFAs, 179 HMM ASVs, 125 gut microbial ASVs at 3 months, 182 gut microbial ASVs at 1 year, and PRSs for asthma and atopy. **(B)** Bar plots show distribution of cases and controls across the two groups of infants. The x-axis shows group membership, and the y-axis shows the percentage of cases (in red) or controls (in green) after excluding the missing values. The controls corresponded to subjects without asthma at age 5 years, no recurrent wheeze between ages 2 to 5 years and no atopic dermatitis (AD) or sensitization to food or inhalant allergens between ages 1 to 5 years. The asterisk symbol (*) corresponds to P values determined by associating group membership with disease prevalence using logistic regression as follows: * P < 0.05 and ** P < 0.01. Also, the tilde symbol (∼) indicates nominal P value significance (P < 0.1).

Our logistic regression analysis also revealed that group membership in the Human Milk-Gut Microbiota-PRS network was significantly associated with the prevalence of asthma and atopy outcomes. For example, Group 2 was associated with reduced prevalence of asthma and atopic outcomes including inhalant sensitization (P=7.0E-3, β=-3.0), recurrent wheeze (P=0.03, β=-1.7), asthma or recurrent wheeze (P=0.02, β=-1.7), atopic dermatitis or sensitization (P=0.02, β=-1.6), or any asthma/atopy outcome (P=0.02, β=-1.3).

## Discussion

Our investigation identified bioactive components in human milk associated with the gut microbiota of milk-fed infants, which in turn may influence the risk of childhood asthma and atopy. Variations in maternal HMOs, HMFAs, and HMM were associated with gut microbial shifts among infants during the first year of life that were previously associated with asthma and atopy by age 5 years^47^. We also demonstrated that host genomics may play an important role in this relationship between exposure to human milk and the gut microbiota, by identifying interactions between HMCs and infants’ PRSs for asthma/atopy. Moreover, we integrated HMCs, infant gut microbial composition, and infant PRSs to cluster infants into different groups associated with disease prevalence. Overall, our study determined interaction effects between HMCs and infant genomics on the infant gut microbiota, which in turn may modulate childhood atopic health and disease susceptibility.

### HMCs may impact gut microbiota in childhood asthma and atopy

Our findings support previous research suggesting that early-life exposure to human milk helps to establish the infant gut microbiota in the first year of life^24,38–43^, a critical period that contributes to long-term risk of childhood diseases^50–52^. We showed that exposure to variable concentrations of HMOs, HMFAs, and HMM may influence gut microbes previously associated with childhood asthma and atopy. For example, exposure to the HMO LNFP3 was associated with decreased levels of the disease-elevated *red* gut microbial cluster, suggesting that HMOs act as beneficial prebiotics for the infant gut microbiota^53–55^. This is supported by an earlier investigation, which reported that increased LNFP3 exposure was associated with reduced cow’s milk allergy^56^. Therefore, our results highlight a potential underlying mechanism by which HMOs may impact infant health and disease through modifications to the infant gut microbial composition.

In contrast to HMOs, few studies have explored the potential impact of HMFAs on the infant gut microbiota and risk of childhood diseases such as asthma and atopy^57^. Exposure to PUFAs such as the n-6 PUFA LA was correlated with disease-elevated microbial features (e.g., *Clostridium sensu stricto sp. 4* and the *yellow* gut cluster). Interestingly, previous studies reported that childhood dietary n-6 PUFAs were associated with asthma^58^, eczema, or sensitization^59,60^. However, other investigations found no associations between n-6 PUFAs in human milk and asthma and other atopic outcomes^57^. Thus, our findings provide novel insights into the relationship between HMFAs and childhood asthma/atopy, suggesting the gut microbiota as a potential mediator, which may explain previously reported discrepancies in the literature.

Our results support the paradigm that HMM plays a critical role in colonizing the infant gut microbiota^39,40,43^. Our study provides unique perspectives, while past investigations were limited to common ASVs shared between the milk and gut microbiota^39^. In contrast, our investigation identified associations between HMM of mothers and microbial network clusters implicated with risk of childhood asthma and atopy. For example, the HMM *Lactobacillus sp. 5* was associated with the disease-reduced *turquoise* gut microbial cluster, whereas the HMM *Rothia sp. 2* was associated with the disease-reduced *salmon* gut microbial cluster. Interestingly, *Lactobacillus* in human milk^61^ or through supplementation^62,63^ is a well-known beneficial probiotic that has been shown to protect against asthma^64–66^. In contrast, reduced gut *Rothia* in early life was previously correlated with increased asthma risk^67^. Therefore, our findings highlight potential mechanisms by which HMM may support the colonization of microbial communities within the infant gut that might protect from asthma and atopy.

### HMCs may interact with infant PRSs to influence their gut microbiota and risk of childhood asthma and atopy

Our G×E interaction analyses provide novel insights into how early-life exposure to HMCs may interact with infants’ PRSs to modify the gut microbiota and risk of asthma and atopy. To the best of our knowledge, we are the first to explore how HMCs may impact the infant gut microbiota depending on host genomics. Our findings demonstrate that exposure to HMOs, HMFAs, and HMM were associated with varying abundances of gut ASVs and microbial network clusters, depending on the infants’ PRSs for asthma or atopy. For example, without considering PRSs, lauric acid in human milk was associated with reduced levels of the disease-elevated *yellow* gut cluster. However, this association was driven by those with high asthma PRS but was not observed in children with low PRS. This demonstrates the importance of considering host genomics for a more nuanced understanding of the relationship between HMCs and the infant gut microbiota. Interestingly, lauric acid was previously shown to reduce tracheal hyper-responsiveness in murine models^68^.

Moreover, G×E interactions identified HMCs associated with the gut microbiota, which may have been missed in earlier studies that did not factor in host genomics. For example, a novel association only detected through G×E analyses included between HMM *P. oryzihabitans* exposure and the disease-elevated *black* gut microbial cluster that was dependent on infants’ PRS. Interestingly, early work from our team found that the presence of *P. oryzihabitans* in human milk was associated with increased asthma prevalence^20^. Thus, our findings highlight a potential mechanism through which exposure to this HMM ASV may impact childhood health, through modulation of the gut microbiota in a genomics-dependent manner. Other investigations also found increased skin *Pseudomonas* was associated with childhood IgE-mediated food allergy^69^ and gut *Pseudomonas* was associated with fecal IgE levels, which are a potential biomarker of childhood dust mite allergies^70^.

Overall, our results suggest that exposure to HMCs could influence the infant gut microbiota and childhood disease risk, depending on the genetic predisposition of the host. This could inform strategies for precision intervention during early life such as supplementation with certain beneficial HMFAs (e.g., lauric acid), particularly among infants with high genetic susceptibility for asthma and atopy, or those who do not receive human milk.

### HMCs, gut microbial composition, and host genomics may jointly identify infants at high risk of childhood asthma and atopy

Past investigations have focused on the impact of individual HMCs^19,20,29–32^ or gut microbial features^71^ on childhood health outcomes. In contrast, through multi-omics subject network analysis we integrated multiple HMCs, infant gut microbiota throughout infancy, and infants’ PRSs to identify groups of infants with differences in asthma and atopy prevalence. The same model without host PRSs was unable to cluster infants into groups associated with disease prevalence. Thus, our findings support the notion that no single HMC, gut microbe, or genomic factor alone determines disease susceptibility. Instead there is a complex interplay among myriad of HMCs, gut microbiota, and genomic factors may jointly influence disease risk. We also demonstrate the importance of utilizing a multi-omics approach for future investigations of childhood asthma, atopy, and other health outcomes.

### Limitations and future directions

While our investigation is the first to integrate infant genomics to study the effects of HMCs on infant gut microbiota and asthma/atopy risk, it is not without limitations. Our investigation utilized stool samples collected at two timepoints in the first year of life. Due to the ever-changing nature of the gut microbiota, especially during early life, more frequent longitudinal stool sample collection throughout infancy and beyond would provide additional insights into how HMCs may shape the composition of the infant gut microbiota. Human milk samples were collected at one timepoint, however, repeated sampling would also allow for studying dynamic changes in milk composition throughout breastfeeding. Thus, future investigations are warranted to explore the relationship between longitudinal changes in human milk exposure and gut microbial composition. Next, additional HMCs (e.g., metabolites, proteins, hormones) not included in our study, may also help to establish the infant gut microbiota and should be investigated in future studies. Additionally, alternative sources of microbiota (i.e. maternal skin and breast pumps), which we were unable to differentiate from HMM, could be ingested and play a role in this relationship. Moreover, future studies using shotgun metagenomics sequencing are necessary to validate our findings, as both gut and milk microbiota data were determined using 16S rRNA marker gene sequencing which restricts our ability to differentiate between lower taxonomic ranks (i.e. species). Lastly, replication of our findings in additional cohorts are necessary but we have yet to identify another cohort with the same combination of HMCs, gut microbiota, and host genomics data.

## Conclusions

Our investigation identified novel associations between exposure to HMCs and infant gut microbial composition by age 1 year, through exploration of microbial ASVs and communities of microbes implicated in childhood asthma and atopy. Moreover, infant genomics may play a crucial role in mediating the association between HMCs and gut microbiota, demonstrating the importance of considering host genomics. Overall, an improved understanding of how HMCs may influence the infant gut microbiota and risk of childhood asthma and atopy will improve our understanding of the mechanisms underlying disease pathogenesis, enable the identification of early-life disease biomarkers, and facilitate the development of targeted prevention strategies– such as supplementation with HMCs–particularly for children with high PRSs or those not receiving human milk.

## STAR ⍰ Methods

### Resource Availability

#### Lead Contact

Further requests for information should be directed to and will be fulfilled by the lead contact, Qingling Duan.

#### Materials Availability

This study did not generate new unique reagents.

#### Data and Code Availability

- The HMM data has been previously deposited into the Sequence Read Archive (SRA) under the BioProject accession numbers (NCBI) PRJNA481046 and PRJNA597997 with SRA number SRP153543.
- The 16S rRNA gut microbiota data has been previously deposited into the SRA under the BioProject accession number PRJNA657821.
- This paper does not report original code.
- Any additional information required to reanalyze the data reported in this paper is available from the lead contact upon request.

### Experimental Model and Subject Details

#### CHILD Cohort Study participants

The CHILD Cohort Study is a prospective, longitudinal Canadian birth cohort that recruited 3,624 women with singleton pregnancies from 2008-2012 from Vancouver, Edmonton, Manitoba, and Toronto (**Figure 1**)^72^. Infant eligibility criteria included birth gestational age ≥ 35 weeks and birth weight ≥ 2,500 grams (N=3,542)^72^. This investigation focuses on subsets of mother-infant dyads with HMOs, HMFAs, or HMM quantified from maternal milk samples collected at 3-4 months postpartum^19–21,23,30,39,73,74^ and infant gut microbiome profiles determined from stool samples collected at 3 months or 1 year of age^39,71,75^ (**Figure 1, Table S1, Table S2**). Genomics data determined from cord blood samples collected at birth was also available for subsets of infants with maternal HMC data and gut microbiota at 3 months or 1 year (**Figure 1**). Additional health, demographic, and exposure information were obtained from physician diagnosis, repeated questionnaires, or hospital records and health care databases, such as childhood asthma and atopy outcomes by age 5 years, infant reported ethnicity, infant sex, and breastfeeding duration (**Table S1**, **Table S6**)^72^. The Human Research Ethics Boards at Queen’s University, McMaster University, the Universities of Manitoba, Alberta and British Columbia, and the Hospital for Sick Children approved study protocols.

#### Childhood asthma and atopy outcomes

Children were assessed for asthma and recurrent wheeze, as well as AD and sensitization to common food or inhalant allergens (**Table S6**)^72^. Asthma at age 5 years was determined by physician clinical assessment by an allergist or asthma specialist and symptom history^72^. Recurrent wheeze between ages 2 to 5 years was defined as ≥ 2 episodes of wheeze in a year without illness, and explored in addition to asthma diagnosis as ‘asthma’ is a heterogeneous label in young children^76,77^. Recurrent wheeze was determined based on extensive repeated questionnaires regarding wheeze triggers and frequency, validated in the International Study of Asthma and Allergies in Childhood^78^. Additionally, AD diagnosis at 1, 3, and 5 years of age was determined by physician assessment and symptom history consistent with the British Association of Dermatologists diagnostic criteria^79^. Skin prick tests (SPT) to common food and inhalant allergens were administered at 1, 3, and 5 years to assess atopic sensitization. Children were considered to sensitized if they had ≥ 1 positive SPTs with wheal diameters ≥ 2mm ^80^. For this investigation, we focused on asthma and atopic outcomes beyond age 1 as earlier symptoms are more likely to be transient during this time^81^.

### Method Details

#### Infant stool sample collection and gut microbiome profiling

Infant stool samples were collected at two timepoints: 3 months (N=779) and 1 year (N=770), and underwent targeted sequencing of the 16S ribonucleic acid V4 hypervariable region, as previously outlined^39,75^. Briefly, data was preprocessed using the DADA2 (v.1.10.0)^82^ pipeline in QIIME2 (v.2019.10)^83^ and the resulting amplicon sequence variants (ASVs) were assigned taxonomic annotations based on 99% similarity using the SILVA database (v.138)^84^. Data filtering was performed to remove ASVs present in less than 5% of subjects, resulting in 125 ASVs at 3 months and 183 ASVs at 1 year^71^. ASV relative abundances were determined, followed by Bayesian-multiplicative zero replacement using the zCompositions R package (v.1.4.0.1)^85^ and centered log-ratio (CLR) transformation using the CoDaSeq package (v.0.99.5)^71,86^. Weighted Correlation Network Analysis (WGCNA) R package^87^ was previously used to identify network clusters of co-occurring gut microbes at 3 months and 1 year, as described^71^.

#### Human milk sample collection and HMC profiling

Human milk samples were collected at 3-4 months postpartum during home visits^21^, and HMOs (n=1206), HMFAs (n=1200), and HMM (n=885) were quantified utilizing various methods. Briefly, 19 of the most abundantly observed HMOs were quantified using high-performance liquid chromatography (**Table S2**)^19,21,73^. These HMOs were classified into four groups: fucosylated/non-sialyated (Fuc+/Sia-), non-fucosylated/sialyated (Fuc-/Sia+), non-fucosylated/non-sialyated (Fuc-/Sia-), and fucosylated/sialyated (Fuc+/Sia+). Additionally, total HMO-bound fucose or sialic acid were determined through summation of total fucose or sialic acid residues concentrations, respectively, while total HMOs were determined through summation of all 19 HMOs concentrations measured^19^.

In addition, 28 HMFA proportions were quantified by gas liquid chromatography and grouped in one or more of the following categories: saturated fatty acids (SFAs), mono-unsaturated fatty acids (MUFAs), poly-unsaturated fatty acids (PUFAs), n-3 PUFAs, and n-6 PUFAs (**Table S2**)^23,30,73^. Total proportions of each of these HMFA categories were computed, in addition to HMFA ratios such as: arachidonic acid (ARA) to docosahexaenoic acid (DHA) ratio (ARA:DHA) and ARA to eicosapentaenoic acid (EPA) and DHA ratio (ARA:EPA&DHA). Both HMO and HMFA concentrations were normalized using rank-based inverse normal transformation from the RNOmni R package (v1.0.1)^88^.

Similar to the gut microbiota, HMM were also determined through targeted sequencing of the 16S ribonucleic acid V4 hypervariable region^39,73,74^, preprocessing with the DADA2^82^ pipeline in QIIME2^83^, and ASVs were identified based on 99% sequencing similarity using the SILVA database^84^. After filtering, 179 ASVs in human milk with a prevalence > 5% remained, followed by the same zero-replacement and CLR transformation as the gut ASVs (**Table S2**).

### Infant genomics data and polygenic risk score computation

Cord blood samples were collected from 2,967 infants at birth^19,71^ for DNA extraction and genotyping of 557,006 single nucleotide variants (SNVs) using the Illumina HumanCoreExome BeadChip. Quality control (QC) using PLINK (v.1.9)^89–91^ and imputations using the Michigan server and Haplotype Reference Consortium data (HRC r1.1 2016) were previously completed^92^. This resulted in 2,835 infants with genomics data and 28 million SNVs, 5.5 million of which were common single nucleotide polymorphisms (SNPs) with a minor allele frequency > 0.05.

This genomics data was used to compute two polygenic risk scores (PRSs), one for asthma or one for atopy, which represent the weighted cumulative genetic risk based on SNPs previously reported in larger population studies. The PRSs were computed using an extension of the Bayesian polygenic prediction method PRS-continuous shrinkage (CS)^93^, called PRS-CSx^94,95^, leveraging PLINK (v.1.9)^89,90^. The PRS-CSx integrates GWAS summary statistics with linkage disequilibrium reference panels from different populations to improve cross-population PRS computation and utilized shared CS priors across populations to estimate posterior SNP effect sizes^94^. We utilized summary statistics from previously published genome-wide association study (GWAS) of childhood-onset asthma by (Pividori *et al.*)^48^ and meta-GWAS of atopic outcomes including asthma, hay fever, and eczema (by Ferreira *et al.*)^49^ as our discovery cohorts. We utilized pre-calculated LD reference panels from the UK Biobank, matched to the ancestry of the discovery GWASs. Moreover population-specific posterior effect size estimates were combined utilizing inverse-variance-weighted meta-analysis using the PRS-CSx “meta” option. The asthma and atopy PRSs were computed based on the cumulative, weighted effect of 1,107,307 and 1,097,013 SNPs, respectively. The resulting PRSs were both z-score normalized.

Logistic regression models in R were used to determine associations between prevalence of childhood asthma and atopy outcomes and our asthma and atopy PRSs, respectively. For our association of asthma PRS with childhood asthma, we utilized two outcomes (1) asthma diagnosis by age 5 years and (2) recurrent wheeze between ages 2 to 5 years. For our association of atopy PRS with childhood atopy outcomes, we utilized four outcomes: (1) atopic dermatitis, (2) food sensitization, (3) inhalant sensitization, and (4) food or inhalant sensitization, all between ages 3 to 5 years. Model covariates included infant sex and the top 10 principal components (PCs), which capture variations in population substructure derived from genomics data, such as genetic ancestry as previously described^19,47^.

### Statistical analysis

#### Association of infant gut microbiota with human milk exposure and polygenic disease risk

Linear regression models in R were used to explore associations between gut microbiota at 3 months and 1 year, and exposure to HMC (HMOs, HMFAs, and HMM) and infants polygenic disease risk. This was achieved through investigating: (i) the main effects of HMC exposure (E) on the infant gut microbiota, (ii) the main effects of infant asthma/atopy PRSs on the infant gut microbiota (G), or (iii) the gene-by-environment (G×E) interaction effects of HMC exposure (E) and infant asthma/atopy PRSs (G) on the infant gut microbiota (**Figure S1**). We focused on 5 gut microbial ASVs (2 at 3 months and 3 at 1 year) and 11 gut microbial network clusters (4 at 3 months and 7 at 1 year) previously associated with childhood asthma and atopy^71^ (**Table S3, Table S4, Table S5**).

Covariates included infant sex, reported ethnicity (White vs. Non-White, derived from parental questionnaires), exact age at stool sample collection (days), time difference between stool/milk sample collection and processing (in seconds), human milk sample processing batch, Study Center (i.e. Vancouver, Edmonton, Manitoba, and Toronto), antibiotics use in the first year of life, and breastfeeding duration. However, in analyses including PRSs (ii and iii), the top 10 PCs were used in place of reported ethnicity.

Multiple testing correction was applied to adjust P values using the Bonferroni correction method. To account for correlations between HMCs we adjusted for 6 clusters of correlated HMOs, 3 clusters of correlated HMFAs, and 85 independent HMM ASVs, were applicable. The 6 clusters of correlated HMOs were previously identified by Ambalavanan *et al.* using Pearson correlation and hierarchical clustering of all 19 HMOs^19^. The 3 clusters of correlated HMFAs were identified through correlation and clustering of 28 HMFAs, using the same method. These clusters of correlated HMOs and HMFAs differ from the different categories described in **Table S2** as they are based on similarities in HMC concentrations, whereas the categories in **Table S2** are grouped based on structural similarities. Moreover, the 85 independent HMM ASVs were previously identified by Fang *et al.* using matSpDlite^96^, a tool utilized for identification of independence based on eigenvalue variance and thus is able to estimate ASVs whose abundance are less likely to be correlated^97^. In analyses containing both HMCs and gut ASVs, we adjusted for both. For example, in analyses exploring associations between gut ASVs at 3 months and HMO exposure we adjusted for 12 tests to account for 6 independent clusters of HMOs × 2 gut ASVs at 3 months (6 × 2 = 12). However, in analyses containing HMCs and gut microbial network clusters we only adjusted for HMCs, as microbial network clusters are not considered independent of one another other^71^. For example, in analyses exploring associations between gut microbial network clusters and HMO exposure, we adjusted for multiple comparisons using Bonferroni correction based on six independent HMO clusters. See **Figure S3** for an overview of multiple testing correction by analysis.

#### Multi-omics subject network construction, clustering, and association with childhood asthma/atopy

We utilized the Similarity Network Fusion (SNF) Tools R package (v.2.3.1)^46^, a tool developed for integration of multiple different types of biological datasets (e.g., transcriptomics, DNA methylation, metabolomics, microbiome, etc.), to construct a multi-omics subject network utilizing maternal HMC, infant gut microbiota, and infant genomic profiles (**Figure S1**). We constructed two multi-omics subject networks through integration of the human milk, gut, and genomics datasets as follows: (1) one including 19 HMOs, 28 HMFAs, 179 HMM ASVs, 125 gut ASVs at 3 months, and 183 gut ASVs at 1 year to explore subject similarity without genetics and (2) one including 19 HMOs, 28 HMFAs, 179 HMM ASVs, 125 gut ASVs at 3 months, 183 gut ASVs at 1 year, asthma PRS, and atopy PRS to explore subject similarity with genetics.

For each network constructed, as recommended by SNF Tools the raw, un-normalized datasets were utilized. These datasets were normalized to have a mean of 0 and standard deviation of 1 using the recommended standardNormalization function^46^. After normalization, the ComBat function in the sva R package (v. 3.54.0) was utilized to remove batch effects from each of the HMO, HMFA, and HMM datasets^98^. Next, individual subject-similarity networks were constructed for each datatype based on Euclidean distance, followed by iterative integration (or fusion) into a single multi-omics subject network. After each network fusion, unsupervised spectral clustering was performed to identify groups of subjects with similarities in input datasets. Two algorithms integrated in the SNF Tools workflow: eigen-gap and rotation cost, were utilized to estimate the optimal number of clusters for spectral clustering.

Once the two subject networks were constructed and infants were clustered into groups as described above, logistic regression models were employed to identify associations between group membership to each network and prevalence of childhood asthma and atopy outcomes, including asthma diagnosis at 5 years, recurrent wheeze between ages 2 to 5 years, and AD, food sensitization, inhalant sensitization, or food or inhalant sensitization between ages 3 to 5 years. Additionally, we explored the combined outcomes of asthma diagnosis or recurrent wheeze, AD or sensitization, and any of the asthma and atopy outcomes explored in this investigation (**Figure S1**). Covariates included infant sex, reported ethnicity or genetic ancestry (captured by the 10 PCs as described above), exact age at stool sample collection, time difference between stool/milk sample collection and processing, human milk sample processing batch, Study Center, antibiotics use in the first year of life, and breastfeeding duration.

## Acknowledgments

We are grateful to all the CHILD families who took part in this study, and the whole CHILD team, which includes interviewers, nurses, computer and laboratory technicians, clerical workers, research scientists, volunteers, managers, and receptionists. For a list of investigators and enrolling centers visit www.childcohort.ca. Computational analyses were performed on resources and with support provided by the Centre for Advanced Computing (CAC) at Queen’s University in Kingston, Ontario. The CAC is funded by the Canada Foundation for Innovation, the Government of Ontario, and Queen’s University. This study was funded by operating grants from CIHR (MRT-168044, PJT-178390).

