## Supplementary Information for "Human milk components interact with infant genomics to modulate gut microbiota, childhood asthma and atopy"

**Table S1.** Overview of characteristics for subsets of infants from the CHILD Cohort Study^1^ with human milk oligosaccharide (HMO), fatty acid (HMFA), or microbiota (HMM) and gut microbiota data at 3 months and 1 year, related to STAR Methods and Figure 1.

| **Infant Characteristics** | **Infants With Gut Microbiota Profiles & HMO Data** | | **Infants With Gut Microbiota Profiles & HMFA Data** | | **Infants With Gut Microbiota Profiles & HMM Data** | |
| --- | --- | --- | --- | --- | --- | --- |
|  | **3 Months (N=402)** | **1 Year (N=394)** | **3 Months (N=398)** | **1 Year (N=393)** | **3 Months (N=292)** | **1 Year (N=296)** |
| Infant Reported Ethnicity | | | | | | |
| White | 249 (61.9) | 255 (64.7) | 245 (61.6) | 253 (64.4) | 187 (64) | 194 (65.5) |
| Non-White | 153 (38.1) | 139 (35.3) | 153 (38.4) | 140 (35.6) | 105 (36) | 102 (34.5) |
| Infant Sex | | | | | | |
| Females | 153 (38.1) | 157 (39.8) | 149 (37.4) | 156 (39.7) | 112 (38.4) | 115 (38.9) |
| Males | 249 (61.9) | 237 (60.2) | 249 (62.6) | 237 (60.3) | 180 (61.6) | 181 (61.1) |
| Human Milk Batch | | | | | | |
| Batch 1 | 242 (60.2) | 219 (55.6) | 241 (60.6) | 218 (55.5) | 163 (55.8) | 153 (51.7) |
| Batch 2 | 160 (39.8) | 175 (44.4) | 157 (39.4) | 175 (44.5) | 129 (44.2) | 143 (48.9) |
| Study Center | | | | | | |
| Edmonton | 67 (16.7) | 63 (16) | 66 (16.6) | 63 (16) | 53 (18.2) | 48 (16.2) |
| Toronto | 122 (30.3) | 106 (26.9) | 120 (30.2) | 105 (26.7) | 89 (30.5) | 85 (28.7) |
| Vancouver | 123 (30.6) | 116 (29.4) | 124 (31.2) | 117 (29.8) | 85 (29.1) | 86 (29.1) |
| Manitoba* | 90 (22.4) | 109 (27.7) | 88 (22.1) | 108 (27.5) | 65 (22.3) | 77 (26) |
| Exact Age At Stool Collection (Years) | 0.3 (0.1) | 1 (0.1) | 0.3 (0.1) | 1 (0.1) | 0.3 (0.1) | 1 (0.1) |
| Time Between Stool Collection and Sample Processing (Hours) | 20.7 (43.5) | 15.4 (19.9) | 20.7 (43.7) | 15.6 (19.9) | 21.7 (49.5) | 14.7 (18.2) |
| Time Between Milk Collection and Sample Processing (Hours) | 28.9 (138.9) | 23.3 (108.3) | 29.2 (139.6) | 23.4 (108.5) | 29 (138.6) | 23.3 (108.1) |
| Antibiotics Use By Age 1 Year | 94 (23.4) | 90 (22.8) | 93 (23.4) | 89 (22.6) | 66 (22.6) | 61 (20.6) |
| Breastfeeding Duration By Age 1 Year (Months) | 12.6 (5.9) | 12.5 (5.9) | 12.6 (5.9) | 12.5 (5.9) | 12.3 (6) | 12.2 (5.9) |
| Values are N(%) or means ± SDs. Percentages reflect proportions of non-missing data. *Manitoba study center includes Winnipeg, Morden, and Winkler. | | | | | | |

**Table S2.** Overview of HMOs, HMFAs, and HMM included in this investigation, related to STAR Methods**.**

| **Human Milk Component Category** | **Human Milk Components** |
| --- | --- |
| **Human Milk Oligosaccharides (HMOs)** | |
| Fucosylated/Non-Sialyated (Fuc+/Sia-) | 2’-fucosyllactose (2’FL)  3-fucosyllactose (3FL)  Difucosyllactose (DFLac)  Difucosyllacto-N-hexaose (DFLNH)  Difucosyllacto-N-tetrose (DFLNT)  Fucosyllacto-N-hexaose (FLNH)  Lacto-N-fucopentaose-I (LNFP1)  Lacto-N-fucopentaose-II (LNFP2)  Lacto-N-fucopentaose-III (LNFP3) |
| Non-Fucosylated/Sialyated (Fuc-/Sia+) | 3’-sialyllactose (3’SL)  6’-sialyllactose (6’SL)  Disialyllacto-N-hexaose (DSLNH)  Disialyllacto-N-tetraose (DSLNT)  Sialyl-lacto-N-tetraose b (LSTb)  Sialyl-lacto-N-tetraose c (LSTc) |
| Non-Fucosylated/Non-Sialyated  (Fuc-/Sia-) | Lacto-N-hexaose (LNH)  Lacto-N-neotetraose (LNnT)  Lacto-N-tetrose (LNT) |
| Fucosylated/Sialyated (Fuc+/Sia+) | Fucodisialyllacto-N-hexaose (FDSLNH) |
| Totals | Total Fucosylated HMOs  Total Sialylated HMOs  Total HMOs |
| **Human Milk Fatty Acids (HMFAs)** | |
| Saturated Fatty Acids (SFAs) | Capric Acid (10:0)  Lauric Acid (12:0)  Myristic Acid (14:0)  Pentadecylic Acid (15:0)  Palmitic Acid (16:0)  Margaric Acid (17:0)  Stearic Acid (18:0)  Arachidic Acid (20:0)  Lignoceric Acid (24:0) |
| Monounsaturdated Fatty Acids  (MUFAs) | Physeteric Acid (14:1n9)  Palmitoleic Acid (16:1n7)  Oleic Acid (18:1n9)  Nervonic Acid (24:1n9)  Vacennic Acid (18:1c11)  Trans Vacennic Acid (TVA) (18:1t11) |
| Polyunstaurdated Fatty Acids (PUFAs) |  |
| n-3 PUFAs | Alpha-Linoleneic Acid (ALA) (18:3n3)  Eicosatetraenoic Acid (20:4n3)  Eicosapentaeonic Acid (EPA) (20:5n3)  Docosapentanoic Acid (DPA) (22:5n3) Docosahexaenoic Acid (DHA) (22:6n3) |
| n-6 PUFAs | Linoleic Acid (LA) (18:2n6)  Conjugated Linoleic Acid (CLA) (18:2c9, t11)  Gamma-Linoleic Acid (GLA) (C18:3n6)  Eicosadienoic Acid (20:2n6)  Dihomo-Gamma-Linoleneic Acid (DGLA) (20:3n6)  Arachidonic Acid (20:4n6)  Adrenic Acid (22:4n6)  Osbond Acid (22:5n6) |
| Totals | Total SFA  Total MUFA  Total PUFA  Total n3 PUFA  Total n6 PUFA |
| Ratios | ARA : DHA Ratio  ARA : EPA & DHA Ratio  n6 PUFA : n3 PUFA Ratio |
| **Human Milk Microbiota (HMM)** | |
| Amplicon Sequence Variants (ASVs) | 179 HMM ASVs |

**Table S3.** Infant gut microbial amplicon sequence variants (ASVs) and network clusters of co-occurring microbes previous associated with asthma and atopy by age 5 years by Stickley *et al.* ^2^, related to STAR Methods.

| **Microbiota Timepoint** | **Microbial Feature Type** | **Microbial Feature** | ***Prior Disease Association** |
| --- | --- | --- | --- |
| 3 Months | Microbial Abundance | *Clostridium sensu stricto sp. 4* | Disease-Elevated |
| 3 Months | Microbial Abundance | *Enterobacter cloacae* | Disease-Elevated |
| 3 Months | Microbial Network Cluster | *Black* | Disease-Elevated |
| 3 Months | Microbial Network Cluster | *Blue* | Disease-Reduced |
| 3 Months | Microbial Network Cluster | *Brown* | Disease-Elevated |
| 3 Months | Microbial Network Cluster | *Turquoise* | Disease-Reduced |
| 1 Year | Microbial Abundance | *Fusicatenibacter sp. 3* | Disease-Reduced |
| 1 Year | Microbial Abundance | *Megasphaera micronuciformis* | Disease-Elevated |
| 1 Year | Microbial Abundance | *Agathobacter sp. 6* | Disease-Reduced |
| 1 Year | Microbial Network Cluster | *Orange* | Disease-Reduced |
| 1 Year | Microbial Network Cluster | *Green* | Disease-Elevated |
| 1 Year | Microbial Network Cluster | *Magenta* | Disease-Reduced |
| 1 Year | Microbial Network Cluster | *Salmon* | Disease-Reduced |
| 1 Year | Microbial Network Cluster | *Red* | Disease-Elevated |
| 1 Year | Microbial Network Cluster | *Purple* | Disease-Reduced |
| 1 Year | Microbial Network Cluster | *Yellow* | Disease-Elevated |
| *Microbial features previously associated with increased asthma/atopy are labelled as disease-elevated, microbial features previously associated with decreased asthma/atopy are labelled as disease-reduced. | | | |

**Table S4.** ASVs in network clusters of co-occurring microbes in the gut at 3 months of age previous associated with asthma and atopy by age 5 years by Stickley *et al.* ^2^, related to STAR Methods.

| **3 Months** | |
| --- | --- |
| **Cluster** | **ASV** |
| *black* | *Lactobacillus sp. 8* |
|  | *Staphylococcus sp. 1* |
|  | *Streptococcus sp. 1* |
|  | *Rothia sp.* |
|  | *Escherichia coli* |
|  | *Lactobacillus paracasei 2* |
|  | *Streptococcus sp. 2* |
|  | *Corynebacterium propinquum* |
| *blue* | *Bifidobacterium animalis* |
|  | *Eggerthella sp.* |
|  | *Blautia sp. 9* |
|  | *Tyzzerella sp. 3* |
|  | *UBA1819 sp.* |
|  | *Blautia sp. 24* |
|  | *Lachnospiraceae UCG-008 sp. 25* |
|  | *[Ruminococcus] gnavus sp.* |
|  | *Flavonifractor sp.* |
|  | *[Eubacterium] oxidoreducens sp.* |
|  | *Unclassified Ruminococcaceae sp.* |
|  | *Anaerostipes sp. 8* |
|  | *Lachnoclostridium sp. 8* |
|  | *Anaerostipes sp. 13* |
|  | *[Clostridium] innocuum sp.* |
|  | *[Ruminococcus] torques sp. 13* |
|  | *Coprobacillus cateniformis* |
|  | *Lachnoclostridium sp. 15* |
|  | *Lachnoclostridium sp. 17* |
|  | *Hungatella sp. 1* |
| *brown* | *Enterococcus pallens* |
|  | *Clostridium sensu stricto sp. 40* |
|  | *Clostridium sensu stricto sp. 4* |
|  | *Clostridioides difficile* |
|  | *Rothia sp. 3* |
|  | *Clostridium sensu stricto sp.* |
|  | *Actinomyces sp.* |
|  | *Bifidobacterium dentium* |
|  | *Citrobacter freundii* |
|  | *Enterobacter cloacae* |
|  | *Enterococcus faecalis 1* |
|  | *Streptococcus equinus* |
|  | *Clostridium butyricum* |
| *turquoise* | *[Clostridium] spiroforme* |
|  | *Roseburia sp. 6* |
|  | *Blautia obeum* |
|  | *Alistipes sp. 1* |
|  | *Roseburia sp. 17* |
|  | *Oscillibacter sp. 2* |
|  | *Lachnospiraceae UCG-001 sp. 2* |
|  | *Anaerostipes sp. 2* |
|  | *Blautia sp. 18* |
|  | *Terrisporobacter sp. 2* |
|  | *Monoglobus sp.* |
|  | *Unclassified Lachnospiraceae sp. 10* |
|  | *[Ruminococcus] torques sp. 19* |
|  | *Anaerostipes sp.* |
|  | *Roseburia sp. 7* |
|  | *Lachnoclostridium sp.* |
|  | *Akkermansia sp.* |
|  | *Ruminococcus sp. 6* |
|  | *Clostridium sensu stricto sp. 14* |
|  | *Akkermansia sp. 3* |
|  | *Fusicatenibacter sp. 3* |
|  | *Alistipes sp. 11* |
|  | *[Ruminococcus] gnavus sp. 15* |
|  | *Ruminococcus sp. 4* |
|  | *[Eubacterium] hallii sp. 3* |
|  | *Faecalibacterium sp. 13* |
|  | *Faecalibacterium sp. 18* |
|  | *Alistipes sp. 4* |
|  | *Dialister sp. 7* |
|  | *[Eubacterium] hallii sp. 1* |
|  | *Parasutterella sp. 1* |
|  | *Agathobacter sp. 6* |
|  | *Roseburia sp. 8* |
|  | *Romboutsia sp. 1* |
|  | *Bacteroides sp. 6* |
|  | *swine fecal* |
|  | *Clostridium sensu stricto sp. 33* |
|  | *Akkermansia sp. 1* |
|  | *Bacteroides sp. 17* |
|  | *Sellimonas intestinalis* |
|  | *Lactococcus lactis* |

**Table S5.** ASVs in network clusters of co-occurring microbes in the gut at 1 year of age previous associated with asthma and atopy by age 5 years by Stickley *et al.* ^2^, related to STAR Methods.

| **1 Year** | |
| --- | --- |
| **Cluster** | **ASV** |
| *green* | *Bifidobacterium bifidum* |
| *green* | *Veillonella seminalis* |
| *green* | *Clostridium sensu stricto sp. 40* |
| *green* | *Lactobacillus sp. 8* |
| *green* | *Clostridium sensu stricto sp. 4* |
| *green* | *Staphylococcus sp. 1* |
| *green* | *Anaerococcus sp.* |
| *green* | *Parabacteroides sp.* |
| *green* | *Streptococcus sp. 1* |
| *green* | *Clostridium sensu stricto sp.* |
| *green* | *Epulopiscium sp.* |
| *green* | *Bifidobacterium longum* |
| *green* | *Varibaculum anthropi* |
| *green* | *Prevotella sp. 6* |
| *green* | *Bifidobacterium longum 1* |
| *green* | *Escherichia coli* |
| *green* | *Klebsiella oxytoca* |
| *green* | *Enterobacter cloacae* |
| *green* | *Enterococcus faecalis 1* |
| *green* | *Streptococcus equinus* |
| *green* | *Lactobacillus paracasei 2* |
| *magenta* | *Abiotrophia sp.* |
| *magenta* | *Solobacterium sp.* |
| *magenta* | *Streptococcus sp. 5* |
| *magenta* | *Streptococcus sp.* |
| *magenta* | *Gemella sp.* |
| *magenta* | *Actinomyces sp. 3* |
| *magenta* | *Shewanella algae* |
| *magenta* | *Streptococcus sp. 2* |
| *orange* | *[Eubacterium] siraeum sp. 1* |
| *orange* | *Roseburia sp. 6* |
| *orange* | *Blautia obeum* |
| *orange* | *Roseburia sp. 17* |
| *orange* | *Anaerostipes sp. 2* |
| *orange* | *Monoglobus sp.* |
| *orange* | *Lachnospiraceae NK4A136 sp.* |
| *orange* | *Anaerostipes sp.* |
| *orange* | *Roseburia sp. 7* |
| *orange* | *[Eubacterium] xylanophilum sp.* |
| *orange* | *Ruminococcus sp. 6* |
| *orange* | *Fusicatenibacter sp. 3* |
| *orange* | *Lachnoclostridium sp. 12* |
| *orange* | *Ruminococcus sp. 4* |
| *orange* | *Faecalibacterium sp. 13* |
| *orange* | *Faecalibacterium sp. 18* |
| *orange* | *Unclassified [Eubacterium]* |
|  | *coprostanoligenes group sp.* |
| *orange* | *Dialister sp. 7* |
| *orange* | *Agathobacter sp. 6* |
| *orange* | *Roseburia sp. 8* |
| *orange* | *Coprococcus sp. 3* |
| *purple* | *[Ruminococcus] torques sp. 7* |
| *purple* | *Erysipelotrichaceae UCG-003 sp.* |
| *purple* | *Lachnoclostridium sp. 19* |
| *purple* | *Lachnospiraceae UCG-001 sp. 2* |
| *purple* | *Butyricicoccus sp. 2* |
| *purple* | *Coprococcus sp. 7* |
| *purple* | *Collinsella sp. 3* |
| *purple* | *Blautia sp. 18* |
| *purple* | *[Eubacterium] hallii sp. 11* |
| *purple* | *[Ruminococcus] torques sp. 19* |
| *purple* | *Unclassified Ruminococcaceae sp. 7* |
| *purple* | *Subdoligranulum sp. 1* |
| *purple* | *Lachnospiraceae UCG-004 sp. 6* |
| *purple* | *Subdoligranulum sp. 15* |
| *purple* | *[Eubacterium] hallii sp.* |
| *purple* | *Dorea sp. 1* |
| *purple* | *Moryella sp.* |
| *purple* | *Lachnospiraceae NK4A136 sp. 9* |
| *purple* | *[Eubacterium] hallii sp. 3* |
| *purple* | *[Eubacterium] hallii sp. 14* |
| *purple* | *Butyricicoccus sp.* |
| *purple* | *Dorea sp.* |
| *purple* | *[Eubacterium] hallii sp. 1* |
| *purple* | *Faecalibacterium sp. 32* |
| *purple* | *Lachnospiraceae ND3007 sp.* |
| *purple* | *Bifidobacterium catenulatum* |
| *purple* | *Lactococcus lactis* |
| *red* | *Veillonella atypica* |
| *red* | *Prevotella sp. 20* |
| *red* | *Haemophilus sp. 3* |
| *red* | *Megasphaera micronuciformis* |
| *red* | *Veillonella sp. 7* |
| *salmon* | *Eisenbergiella sp. 4* |
| *salmon* | *Intestinimonas butyriciproducens* |
| *salmon* | *Akkermansia sp.* |
| *salmon* | *Alistipes sp. 11* |
| *salmon* | *Gordonibacter faecihominis* |
| *yellow* | *Bifidobacterium animalis* |
| *yellow* | *Eggerthella sp.* |
| *yellow* | *Lachnoclostridium sp.* |
| *yellow* | *Blautia sp. 9* |
| *yellow* | *Eisenbergiella sp.* |
| *yellow* | *Lachnospira sp. 3* |
| *yellow* | *Tyzzerella sp. 3* |
| *yellow* | *Anaerotruncus sp.* |
| *yellow* | *[Ruminococcus] gnavus sp.* |
| *yellow* | *Flavonifractor sp.* |
| *yellow* | *[Eubacterium] oxidoreducens sp.* |
| *yellow* | *Lachnoclostridium sp. 8* |
| *yellow* | *Anaerostipes sp. 13* |
| *yellow* | *[Clostridium] innocuum sp.* |
| *yellow* | *Coprobacillus cateniformis* |
| *yellow* | *Lactobacillus acidophilus* |
| *yellow* | *Lachnoclostridium sp. 15* |
| *yellow* | *Lachnoclostridium sp. 17* |
| *yellow* | *Clostridium sensu stricto sp. 44* |
| *yellow* | *Hungatella sp. 1* |
| *yellow* | *Streptococcus salivarius* |

**Table S6.** Overview of asthma and atopy outcomes by age 5 years explored in this study, related to STAR Methods, Figure 3, Figure 6, and Figure 7.

| **Asthma/Atopy Outcome** | **Infants With Genomics Profiles  (n=2835)** | **Infants With Human Milk Components (HMC) & Gut Microbiota Profiles  (n=218)** | **Infants With HMC, Gut Microbiota & Genetics Profiles  (n=199)** |
| --- | --- | --- | --- |
| Asthma (5Y) | 145 (13.7) | 31 (34.8) | 28 (34.1) |
| Recurrent Wheeze (2-5Y) | 381 (29.4) | 58 (50) | 53 (49.5) |
| Asthma or Recurrent Wheeze | 413 (31.1) | 63 (52.1) | 58 (51.8) |
| Atopic Dermatitis (3-5Y) | 463 (33.6) | 50 (46.3) | 46 (46) |
| Inhalant Sensitization (3-5Y) | 533 (36.9) | 59 (50.4) | 54 (50) |
| Food Sensitization (3-5Y) | 182 (16.6) | 33 (36.3) | 31 (36.5) |
| Food or Inhalant Sensitization | 592 (39.3) | 69 (54.3) | 63 (53.8) |
| Atopic Dermatitis or Sensitization (3-5Y) | 857 (48.4) | 84 (59.2) | 77 (58.8) |
| Any Asthma or Atopy Outcome | 1053 (53.6) | 114 (66.3) | 104 (65.8) |
| Values are N(%) or means ± SDs. Percentages reflect proportions of non-missing data. Controls are subjects without asthma by 5 years of age, recurrent wheeze between ages 2 to 5 years, or atopic dermatitis (AD) or food or inhalant sensitizations between ages 1 to 5 years. | | | |

**Table S7.** Summary of HMOs, HMFAs, and HMM associated (PBonferroni < 0.05) with gut microbiota previously correlated with asthma and atopy, related to STAR Methods and Figure 2. *See excel table.*

**Table S8.** Summary of G×E interactions between infants asthma polygenic risk score and HMOs, HMFAs, and HMM associated (P_Bonferroni_ < 0.05) with gut microbiota previously correlated with asthma and atopy, related to STAR Methods and Figure 4. *See excel table.*

**Table S9.** Summary of G×E interactions between infants atopy polygenic risk score and HMOs, HMFAs, and HMM associated (P_Bonferroni_ < 0.05) with gut microbiota previously correlated with asthma and atopy, related to STAR Methods and Figure 5. *See excel table.*

**Table S10.** Summary of associations between multi-omics subject network group membership to the Human Milk-Gut Microbiota network and asthma and atopy prevalence, related to STAR Methods and Figure S6.

| **Infant Health** | **Human Milk-Gut Microbiota Network** | **P** | **Beta** | **CI 2.5%** | **CI 97.5** |
| --- | --- | --- | --- | --- | --- |
| Inhalant Sensitization | Group 2 Membership | 0.07 | 1.12 | -0.06 | 2.44 |
| Food or Inhalant Sensitization | Group 2 Membership | 0.19 | 0.69 | -0.30 | 1.75 |
| Atopic Dermatitis | Group 2 Membership | 0.22 | 0.72 | -0.39 | 1.92 |
| Atopic Dermatitis or Sensitization | Group 2 Membership | 0.28 | 0.53 | -0.41 | 1.51 |
| Food Sensitization | Group 2 Membership | 0.30 | 0.70 | -0.57 | 2.12 |
| Asthma Diagnosis | Group 2 Membership | 0.38 | 0.64 | -0.74 | 2.14 |
| Recurrent Wheeze | Group 2 Membership | 0.43 | 0.44 | -0.63 | 1.58 |
| Asthma or Recurrent Wheeze | Group 2 Membership | 0.49 | 0.37 | -0.67 | 1.46 |
| Any Outcome | Group 2 Membership | 0.74 | 0.14 | -0.69 | 0.98 |

**Table S11.** Summary of associations between multi-omics subject network group membership to the Human Milk-Gut Microbiota-PRS network and asthma and atopy prevalence, related to STAR Methods and Figure 6.

| **Infant Health** | **Human Milk-Gut Microbiota-PRS Network** | **P** | **Beta** | **CI 2.5%** | **CI 97.5** |
| --- | --- | --- | --- | --- | --- |
| Inhalant Sensitization | Group 2 Membership | 7.04E-03 | -3.02 | -5.64 | -1.12 |
| Any Outcome | Group 2 Membership | 2.05E-02 | -1.28 | -2.41 | -0.23 |
| Atopic Dermatitis or Sensitization | Group 2 Membership | 2.22E-02 | -1.57 | -3.02 | -0.29 |
| Asthma or Recurrent Wheeze | Group 2 Membership | 2.43E-02 | -1.73 | -3.39 | -0.32 |
| Recurrent Wheeze | Group 2 Membership | 2.96E-02 | -1.75 | -3.49 | -0.28 |
| Food or Inhalant Sensitization | Group 2 Membership | 5.21E-02 | -1.36 | -2.84 | -0.05 |
| Atopic Dermatitis | Group 2 Membership | 1.65E-01 | -1.17 | -2.97 | 0.42 |
| Food Sensitization | Group 2 Membership | 7.21E-01 | -0.34 | -2.29 | 1.55 |
| Asthma Diagnosis | Group 2 Membership | 9.99E-01 | -206.99 | -7163.82 | 6749.85 |

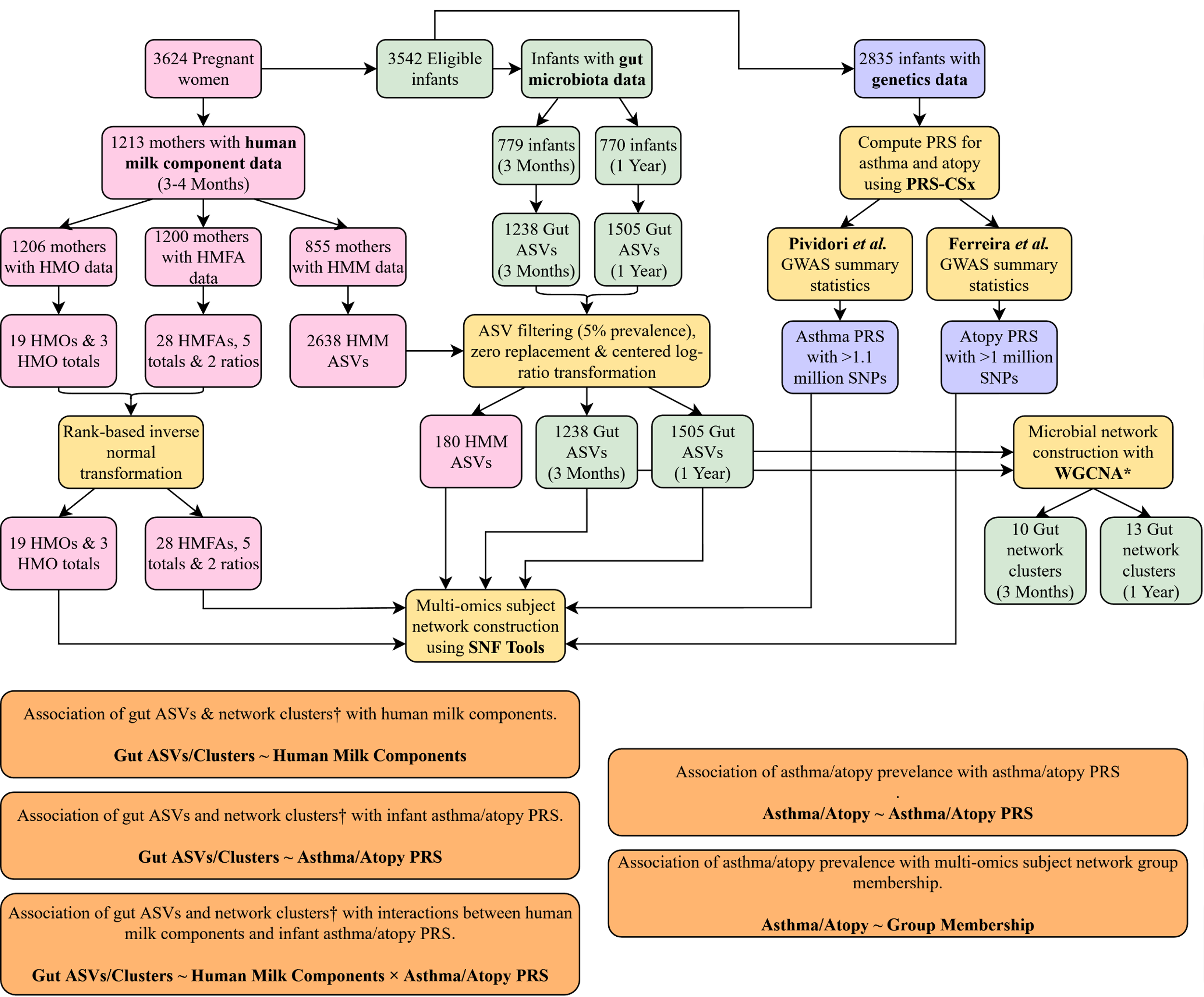

**Figure S1. Overview of study design, related to STAR Methods. ***Microbial network construction with weighted gene correlation network analysis (WGCNA) was performed by Stickley *et al.* previously^2^. † Focused on gut ASVs and network clusters previously associated by Stickley *et al.* with asthma and atopy outcomes by age 5 years. ASV: Amplicon sequence variants; PRS: Polygenic risk score; PRS-CSx: PRS score-continuous shrinkage x; SNF Tools: Similarity Network Fusion Tools

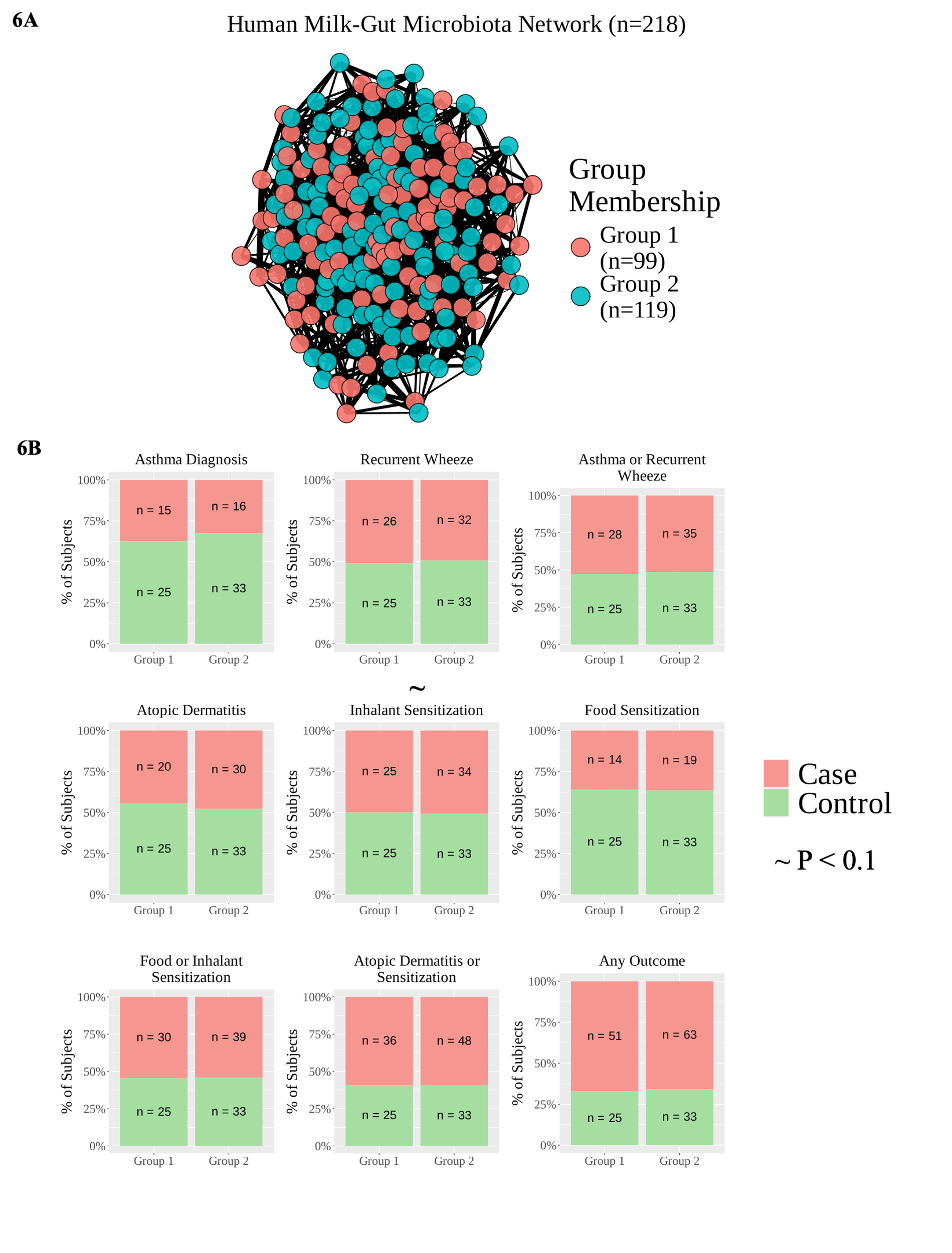

**Figure S2. Multi-omics integration of HMCs and infant gut microbiota. (A)** Network plots show two groups of infants determined from multi-omics subject network construction and clustering of 19 HMOs, 28 HMFAs, 179 HMM ASVs, 125 gut microbial ASVs at 3 months, and 182 gut microbial ASVs at 1 year, without inclusion of asthma/atopy PRSs. **(B)** Bar plots show distribution of cases and controls across the two groups of infants. The x-axis corresponds to the group membership and the y-axis to the percentage of cases (shown in red) or controls (shown in green) after excluding missing values. The controls corresponded to subjects without asthma at age 5 years, no recurrent wheeze between ages 2 to 5 years, and no atopic dermatitis (AD) or sensitization to food or inhalant allergens between ages 1 to 5 years. The tilda symbol (~) indicates nominal P value significance (P < 0.1).

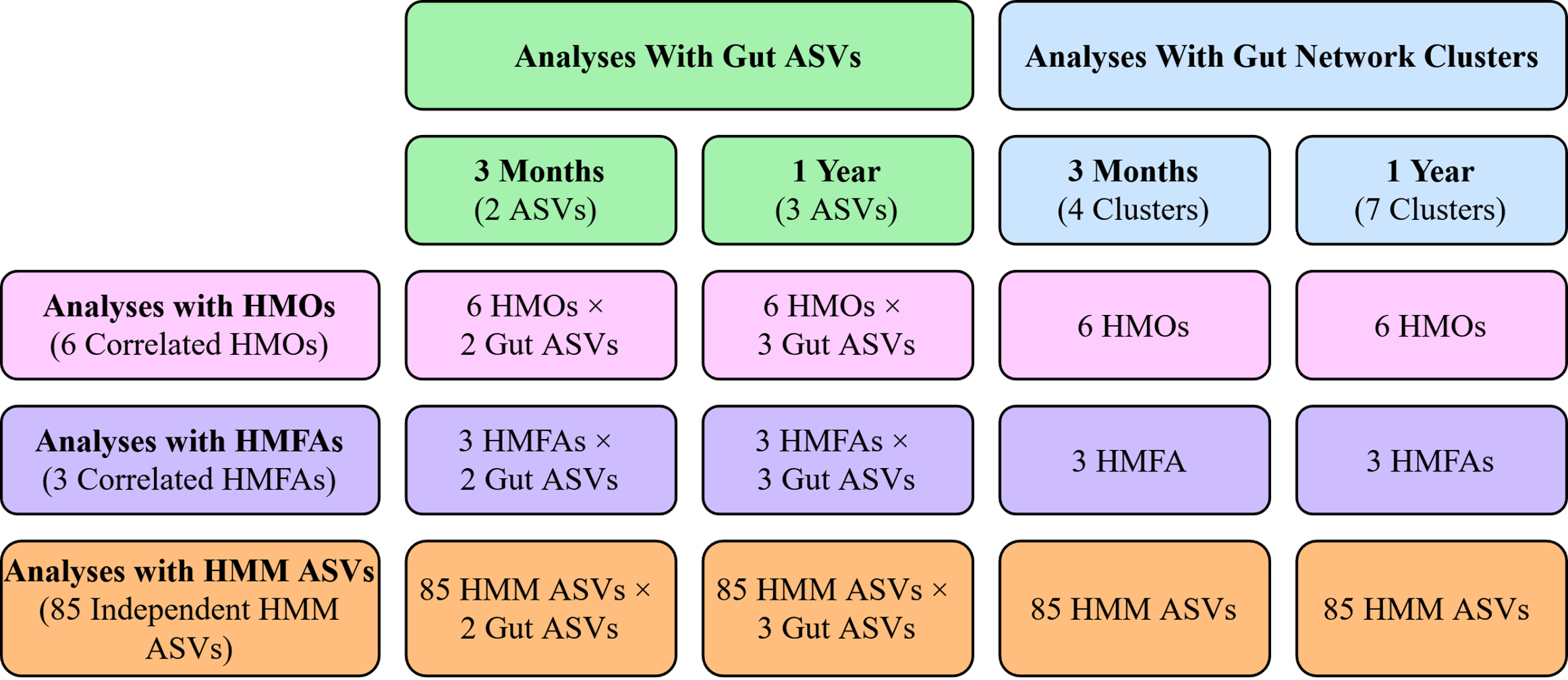

**Figure S3. Overview of multiple testing correction by analysis, related to STAR Methods.** Previous studies by Ambalavanan *et al.*, identified 6 clusters of correlated HMOs through Pearson correlation and hierarchical clustering^3^. Moreover, 3 clusters of correlated HMFAs were identified using the same methods. Notably, these clusters are determined based on similarities in HMC concentrations, instead of structural similarities that dictate the different HMC categories in Table S2. Additionally, previous studies by Fang *et al.* identified 85 independent HMM ASVs based on similarities in abundances^4^.
